# Representations of active grasp maintenance emerge during reach-grasp-carry learning in mouse motor cortex

**DOI:** 10.64898/2026.06.30.735546

**Authors:** Eliza T. Wiener, Gabriella Wheeler Fox, Jason N. MacLean

**Affiliations:** Department of Neurobiology, University of Chicago, Chicago, IL 60637, USA; Committee on Computational Neuroscience, University of Chicago, Chicago, IL 60637, USA

**Keywords:** motor cortex, grasping, learning, functional networks, dexterity, prehension

## Abstract

Object manipulation is a dexterous behavior requiring ongoing sensorimotor coordination to establish and maintain a grasp during consecutive movements. Although grasp initiation is known to modulate activity in primary motor cortex (M1), it remains unclear if sustained grasp maintenance is represented in M1, whether it corresponds to an intrinsic M1 state or one that emerges through learning, and how this state contributes to skilled behaviors involving object manipulation. We addressed these questions using two-photon calcium imaging in mice learning a self-initiated, self-paced, reach-grasp-carry task. This task allowed mice to control trial structure and timing, revealing two kinematically similar but functionally distinct carry behaviors: active grasps, in which mice maintained control of a sucrose pellet and received sensory feedback from the held object, and empty grasps, in which mice completed the same carry movement after failing to grasp the pellet. As behavior was refined, empty grasps became less frequent and the M1 representation of active grasp maintenance became more reliable and more distinct from empty grasp. This distinction was evident in single-neuron activity, population activity, and functional connectivity between neurons. These differences were sufficiently robust to support accurate decoding of active and empty grasp state from neural activity and improved encoding of single neuron activity when models incorporated condition-specific functional network coupling. Together, these findings show that learning refines M1 activity into a distinct and reliable representation of active grasp maintenance, revealing sensorimotor integration during object contact as a key neural substrate of learned dexterous behavior.

## INTRODUCTION

Maintaining a grasp during subsequent movements is not a simple motor behavior; it requires ongoing coordination of digit posture, grip force, and sensorimotor feedback to ensure that the object remains secured. Although grasping has been extensively studied in primates and, to a lesser extent, in rodents ^1–8^, less is known about active grasp maintenance and how it is learned during motor skills that require sustained object control. To fully comprehend how sensorimotor integration underlying object manipulation occurs, we must first understand whether the primary motor cortex (M1) represents the sensorimotor state of sustained grasp–in addition to the kinematics related to grasping–and characterize how these representations emerge.

Although it remains unclear how M1 represents the maintenance of a grip during subsequent movements, M1 is strongly implicated in grasp control. Inactivation or disruption of M1 broadly impairs skilled forelimb movements ^9,10^, and M1 is specifically implicated in controlling hand shape and grip force during grasping ^7,8^. More recent work in primates has characterized separable M1 representations of grasp by grasp type, hand shape, and grip force ^4,5^, while also showing that grasp representations can be higher dimensional than representations of simpler actions such as reaching ^6^. The visual presence of an object, somatosensory stimulation of the hand, and tactile sensory expectation like that experienced during active grasping can also modulate M1 activity and alter population dynamics ^11–16^. This body of work establishes that grasping depends on sensorimotor interactions distributed across cortical circuits. However, most studies of grasp-related activity have focused on grasp initiation, grip type, or force production during discrete grasp-and-release behaviors. Much less is known about how M1 represents the maintenance of an object-related grip during a subsequent movement, such as carrying an object to the mouth.

It is additionally unknown how this representation of maintained grasp might develop over learning. Learning alters M1 activity patterns, dendritic structure, and functional interactions among neurons ^17–22^, and inactivation of M1 can prevent the acquisition of skilled movements ^23^. Examining active grasp maintenance across learning therefore provides an opportunity to determine how a sensorimotor representation emerges as behavior is refined. This question is especially important for dexterous skills: whereas some learned non-dexterous behaviors can be executed with reduced M1 engagement, reach-grasp-carry behavior requires M1 for both learning and movement execution ^8–10^. Thus, characterizing how the representation of active grasp maintenance in M1 develops over learning can reveal how sensorimotor integration is organized during the acquisition and performance of dexterous motor skills.

Here, we investigated how M1 represents active grasp maintenance as mice learned a self-initiated, self-paced, reach-grasp-carry task. Using two-photon calcium imaging, we tracked M1 activity as animals learned to reach for, grasp, and carry a sucrose pellet to the mouth. Because mice controlled trial structure and timing, they were free to continue attempts after failing to secure the pellet. Thus, this task produced two closely related carry behaviors: active grasp, in which mice established and maintained a grip on the pellet during carry, and empty grasp, in which mice performed a similar carry movement without a grasped pellet. This naturalistic structure allowed us to separate the sensorimotor state of active grasps from the kinematics of a more general grasp-carry motion, isolate neural activity associated only with active grasp maintenance, and examine how that representation changed as behavior was refined. We find that, over learning, active grasp maintenance becomes represented as a distinct M1 state expressed in single-neuron activity, population activity, and functional network structure, identifying M1 as a site where object-related sensory feedback is integrated into the control of skilled dexterous behavior.

## RESULTS

### Mice perform self-paced reach-grasp-carry task

Head-fixed mice were trained to reach for a sucrose pellet placed on a narrow pedestal. The task was not externally cued; instead, each trial began at the onset of a self-initiated reach. After a pellet was presented, mice typically made a bout of several reaches. Each bout ended in either success–defined as grasping the pellet, carrying it to their mouth, and consuming it–or failure–in which the pellet was either knocked off of the pedestal or dropped during the carry movement.

The pedestal diameter was similar to that of the sucrose pellet, making the pellet easy to dislodge, and consequently, rendering the task challenging. Because the narrow pedestal limited paw repositioning after contact with the pellet, mice had to execute an accurate reach and grasp from the outset, requiring precise paw placement and coordinated digit control. Successful retrieval then required that they maintain their grasp while transporting the pellet to their mouth, further increasing task complexity. Performance, measured by success rate, plateaued after 2-5 days of training in the majority of mice.

We recorded mouse behavior using two high-speed (200 fps) cameras and employed markerless keypoint tracking to reconstruct the kinematics of each reach ^24,25^. Using keypoint tracking from both cameras, we triangulated keypoint locations to create a three-dimensional rendering of the mouse forelimb, allowing the tracking of detailed kinematic information (Figure 1A). Tracking of the elbow and wrist reflected activation of larger forelimb muscles involved in larger reaching and carrying movements, while tracking of the digits allowed for reconstruction of digit kinematics and paw posture during grasping.

**Figure 1.**
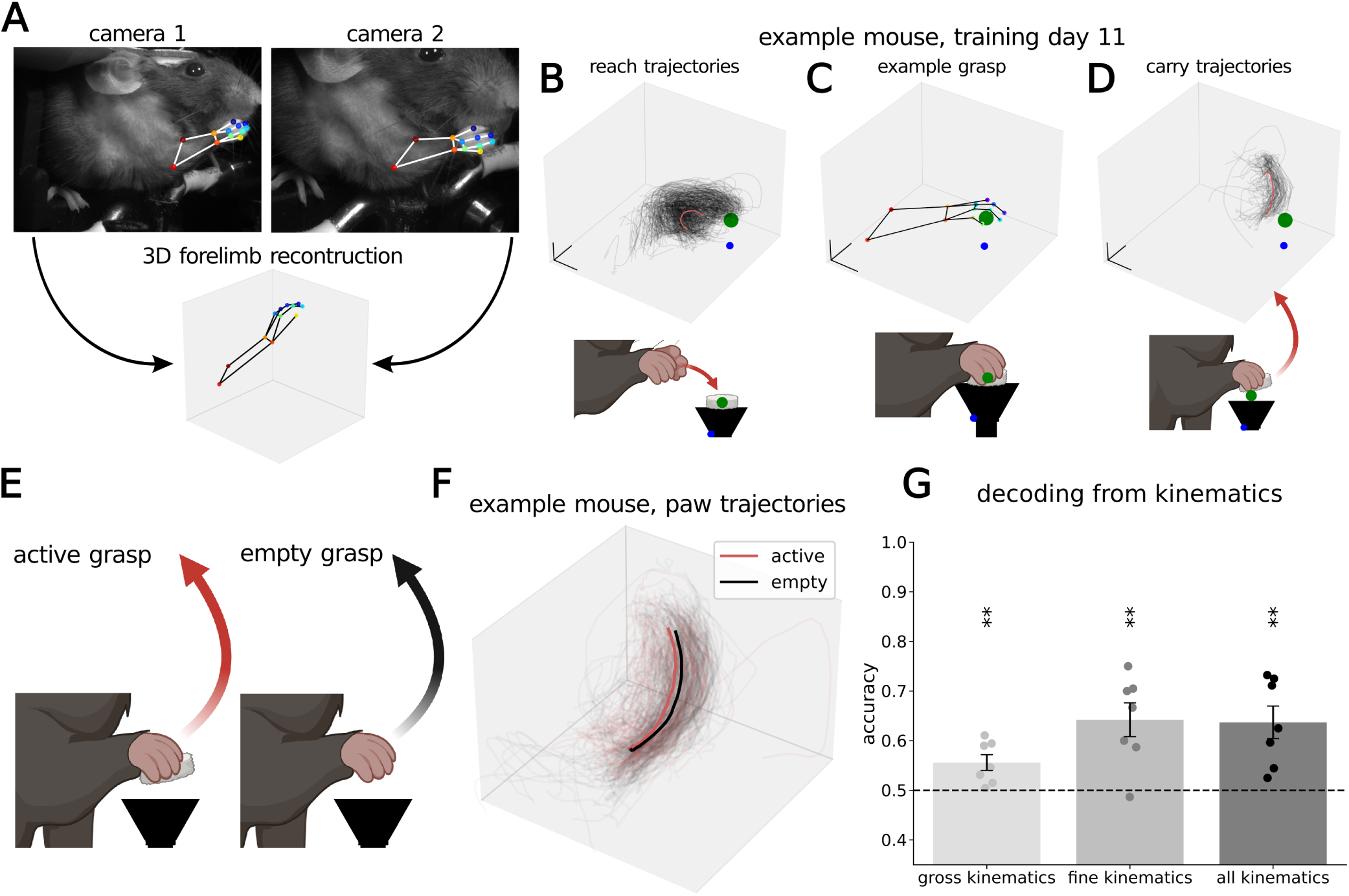
Mice perform reach-grasp-carry task. A) Stills from two cameras and 3D triangulation of paw and forelimb from these perspectives. B-D) (top) Behavioral instances, green point indicates pellet position on the pedestal, blue point indicates pedestal keypoint label, red lines in (B) and (D) indicate mean trajectory. Black axis lines indicate scale. (bottom) Cartoonized graphic of behavioral instance. B) Trajectories of paw centroid during reach, n=324 trials. C) Example grasp still, all keypoints. D) Carry trajectories, n=83 trials. E) Schematic of active and empty grasp. F) Example mouse, training days 3-6. Active and empty grasp paw centroid trajectories. Average trajectory shown in bold lines. Red lines are active grasp, black lines are empty grasp. G) Decoding performance for predicting trial category (active and empty grasp) from kinematics. Accuracy is average across mice, error bars are standard error across mice. Points are cross-validated accuracies for individual mice (n=7 mice). Gross kinematics: p=0.006; fine kinematics: p=0.003; all kinematics p=0.003 by one-sample t-test vs chance of 0.5. \*\**p <* 0.01. Cartoon graphics (B-D bottom, E) created with BioRender.com/a2xeh0g

We segmented the behavior into reach, grasp, and carry phases. The reach phase primarily involved coordinated activation of proximal muscle groups to move the forelimb toward the pedestal (Figure 1B), whereas the grasp phase required distal muscle coordination and the resultant fine digit control to close around the pellet (Figure 1C). The carry phase required coordinated gross and dexterous control to maintain the grasp while transporting the pellet to the mouth (Figure 1D). Because successful task completion depended on maintaining the grasp during carry so that the pellet can be consumed, we focused our analyses on the carry phase of the behavior.

During the carry phase, trials fell into two categories: active grasp and empty grasp. Active grasps were carry instances in which mice established a grip on the pellet and maintained it during paw transport to the mouth (Figure 1E, Video S1). Empty grasps occurred when mice proceeded into the carry phase of the behavior without first establishing a successful grasp of the pellet (Figure 1E, Video S2).

### Learning shifts carry behavior from empty to active grasp

Active and empty grasp trials had nearly indistinguishable large-scale kinematics, with negligible differences in the correlation between keypoint trajectories within or across carry categories (Figure 1F, mean correlation difference = 0.006). Consistent with this similarity, decoding of active or empty trial category from gross kinematics performed only slightly above chance (mean accuracy = 0.56 *±* 0.02; n=7 mice, Figure 1G). Despite the similarity in gross kinematics, the two trial types differed fundamentally in potential behavioral outcome. While active grasps did not always lead to successful pellet retrieval and consumption, with drops occurring in the transition between single paw carry and dual paw eating behaviors, they retain the possibility of sucrose reward throughout the trial. In contrast, empty grasps could never result in reward. By comparing behaviors that are kinematically similar but functionally distinct–carry movements with the presence or absence of an active grasp–we evaluated whether M1 represents the complete sensorimotor state of active grasp maintenance or the movement trajectory alone.

Empty grasps decrease in prevalence over learning, with the exact timing of the decrease varying across animals. On average, the number of empty grasp trials peaked at training day 3 (”empty grasp peak”, Figure 2A) and ranged between training days 1-4 across individual mice (Figure 2C). The number of empty grasp trials then fell, and the proportion of active grasp trials increased to exceed that of empty grasp trials, marking the “active-empty crossover point”. Across mice, this crossover occurred on average on training day 5 (Figure 2B), with individual crossover points ranging from training days 2 to 7 (Figure 2C). In mouse 7, the proportion of active grasps approached, but never exceeded, the proportion of empty grasps. For analysis, we consider the crossover point for mouse 7 to be training day 7, when active and empty grasp proportions stabilized.

**Figure 2.**
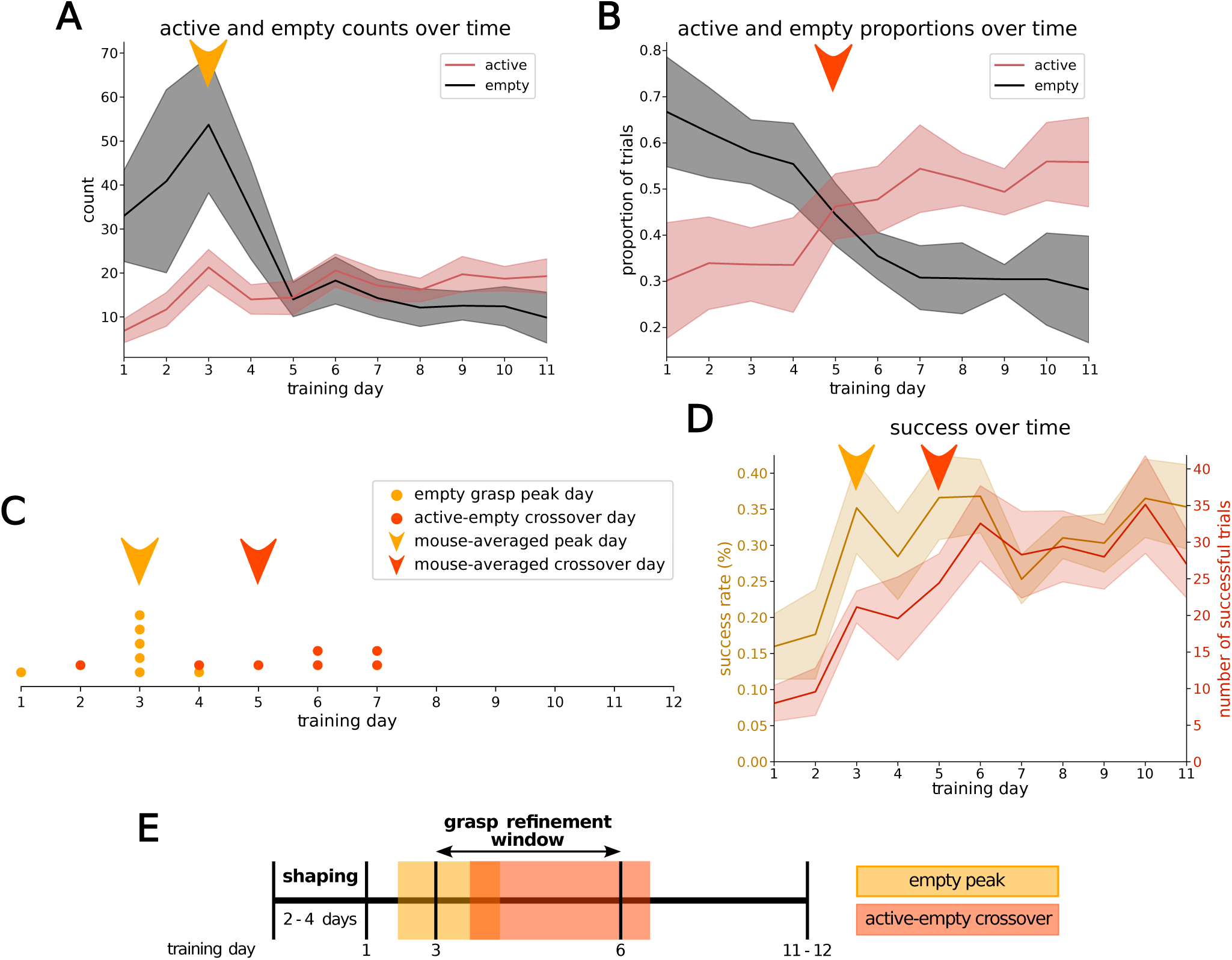
Behavioral shift from empty grasp to active grasp defines a grasp refinement period. A) Number of trials by category. Active grasp in red, empty grasp in black. Lines are average count over mice, shaded region around lines is standard error across mice. Yellow arrow marks mouse-average empty grasp peak (n=7 mice). B) Proportion of active (red) and empty (black) carry trials by category over learning. Lines are average proportions across all mice, shaded region is standard error across mice. Orange arrow marks mouse-averaged active-empty crossover point (n=7 mice). C) Timeline of empty grasp peak and active-empty crossover. Each point represents the peak (yellow) or crossover (orange) day for a single mouse. Arrows indicate mouse-average peak (yellow) or crossover (orange) day. D) Task performance over learning. Yellow indicates success rate. Red-orange line indicates number of successes (pellet consumptions). Lines are average across mice, shaded area is standard error across mice. Yellow arrow marks mouse-average empty grasp peak. Orange arrow marks mouse-averaged active-empty crossover point (n=7 mice). E) Schematic of training timeline and grasp refinement period. Yellow box is range of empty peak days across mice. Orange box is range of active-empty.

Behaviorally, the empty grasp peak and active-empty crossover defined the bounds of a “grasp refinement window” during learning, which generally occurred between training days 3 and 6 (Figure 2F). At the beginning of the window, marked by the empty grasp peak, animals were performing many empty grasp trials and any given carry instance was considerably more likely to be an empty grasp than an active grasp (Figure 2A, C, E). During the grasp refinement window, the number of empty carries decreased because mice were less likely to complete the carry movement after failing to grasp the pellet, instead retracting their paw to the handlebar, which allowed for the delivery of a new pellet. The crossover point marked the end of the window, when animals became more likely to perform active grasps than empty grasps (Figure 2B-C, E).

This refinement affected the number of successful trials without producing a comparably large change in success rate. On average across mice, success rate plateaued by training day 3, whereas the number of successful trials continued to rise over the grasp refinement window and did not plateau until training day 6 (Figure 2D). This distinction arises because success rate measures the fraction of pellet presentations that result in success and therefore does not capture repeated empty grasps that occur after a pellet has already been dislodged from the pedestal. Increasing the total number of successful trials required mice to refine their behavior so that they aborted trials after the reward was no longer available, thereby allowing more attempts with the possibility of success. Improved reward consumption in this reach–grasp–carry task therefore required multiple behavioral refinements, including recognition of pellet presence and improvement of grasp posture and grip force. Correspondingly, decoding trial category from fine-scale paw kinematics, including aperture, splay, and orientation, performs modestly but significantly better than chance (mean accuracy = 0.64 *±* 0.03; one sided t-test, p=.0029, Figure 1G). The fine kinematic decoder indicates that active grasp maintenance in part corresponds to subtle paw postural changes as compared to empty grasp. Together, these results suggest that learning may depend in part on increasing differentiation between empty or improper grasp postures and active grasp maintenance, potentially through improved use of sensory feedback during grasp and carry.

### Active grasp maintenance modulates population and single cell activity

Behavioral refinements over learning could be supported by differences in neural activity encoding the sensorimotor changes between active and empty grasp. We next sought to characterize these neural representations underlying the distinction between active and empty grasp. To obtain sufficient trial counts for robust decoding in all mice, we pooled trials across the grasp refinement window and restricted analyses to simultaneously recorded cells that were present on all days (Figure 2D; training days 3-6 for mice 2-7, training days 2-6 for mouse 1).

Despite their kinematic and behavioral similarity, active and empty grasp trials were clearly differentiated by population and single cell activity. We first tested whether active and empty grasps could be decoded from single-neuron activity. For each neuron, we trained two support vector machine (SVM) models: one using a single feature, the time-averaged firing rate across the trial, and a second using eight features, each corresponding to the instantaneous firing rate in a single bin. A neuron was classified as ‘decodable’ if at least one model achieved cross-validated accuracy with 95% confidence interval above chance. Just under half of all neurons (48.6%, n=1209/2486) were decodable (Figure 3D). Decodable neurons exhibited diverse response profiles. Some showed higher firing rates during active grasp than empty grasp (n=522, example in Figure 3A) and some showed lower firing rates during active grasp (n=433, example in Figure 3B). A smaller subset was decodable only when time varying dynamics were included in the instantaneous firing-rate model (n=204, example in Figure 3C). This diversity, including neurons positively modulated by one condition or the other and neurons with temporally structured differential activity, indicates robust single cell encoding of active grasp maintenance.

**Figure 3.**
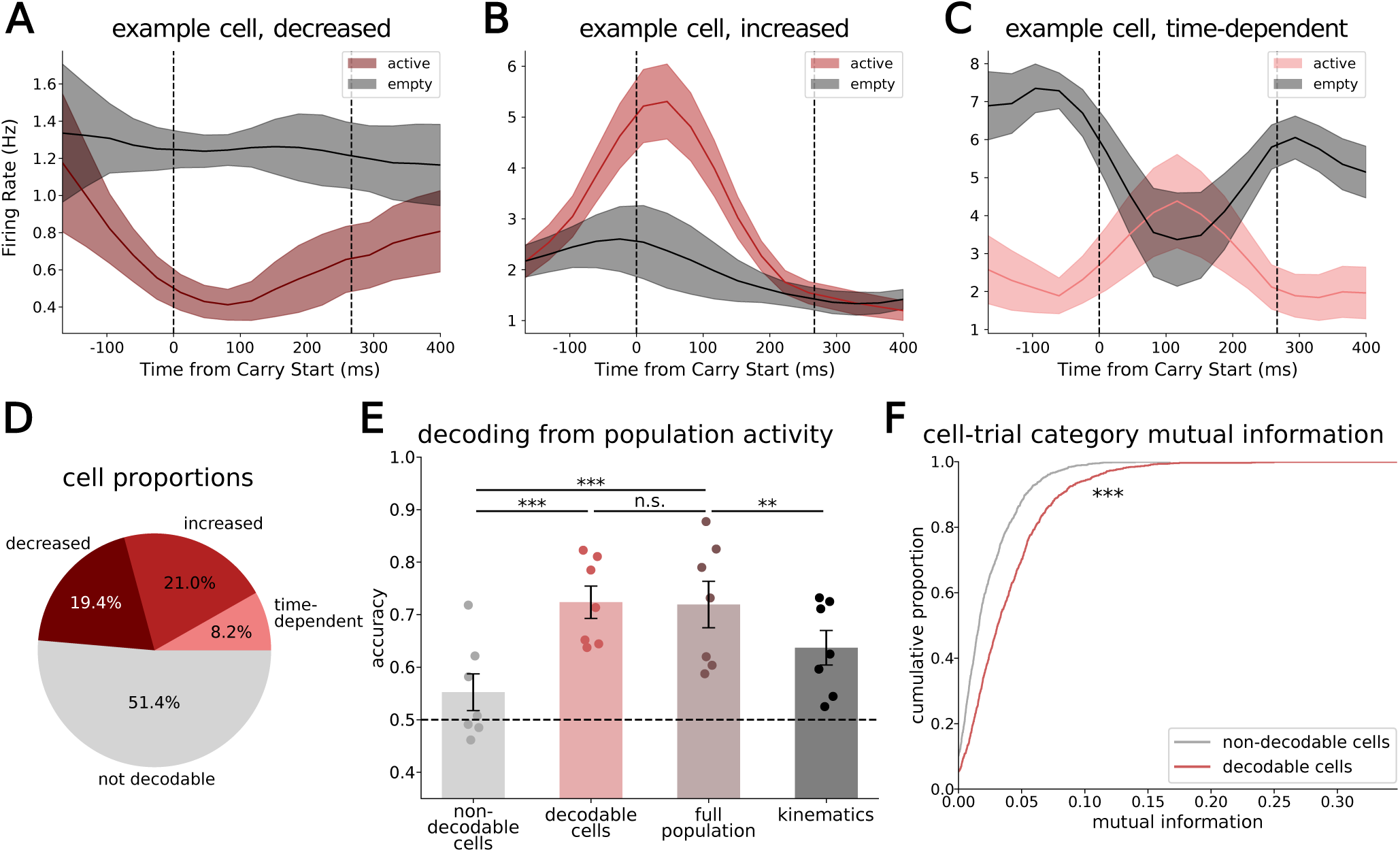
Active grasp maintenance modulates population and single cell activity. A-C) Example decodable cells with increased firing rate (A), decreased firing rate, (B), and time-varying modulation (C). Lines are average across trials, shaded areas are standard error across trials. Black dotted lines indicate start and end of carry trial. D) Proportions of cells by decodability and type of decodable neuron. E) Decoding performance by cell population. Accuracy is average across mice, error bars are standard error across mice. Points are individual mouse cross-validated accuracies. Non-decodable cell decoding accuracy was lower than decodable cells (one-sided paired t-test by mice, *p* = 1.9 *∗* 10*^−^*^4^) and full population (one-sided paired t-test by mice, *p* = 3.5 *∗* 10*^−^*^4^). Decodable cells and full population decoded similarly (two-sided paired t-test by mice, *p* = .773). Full population decoding accuracy was higher than kinematics (two-sided paired t-test by mice; n=7 mice). F) Cumulative distribution of mutual information between individual neurons and trial category for decodable (red, n=1209 cells) and non-decodable (grey, n=1277 cells) neurons. *p <* 10*^−^*^16^ by Wilcoxon rank-sum. *** *p <* 0.001; n.s. *p* > .05. Also see Figure S1.

Trial type could also be decoded from the full population activity with high accuracy (mean accuracy = 0.72*±*0.04; n=7 mice) (Figure 3E). This performance was significantly higher than decoding from gross, fine, or combined kinematics (Figures 3E, S1A), indicating that neural activity can access information that distinguishes between active and empty grasp beyond the paw position, shape, and orientation included in kinematic decoders. We then asked how different neural subpopulations contributed to population decoding. While non-decodable neurons could not individually distinguish between active and empty grasp with linear decoding, it is possible that they represent active-empty grasp differences at a population level. Although decodable neurons account for most of the population decoding (decoding accuracy similar to that of the full population; paired t-test, *p* = .773), non-decodable subpopulations supported limited but above-chance decoding of trial type in some mice (Figure 3E). Looking at the mutual information between single neural activity and trial category, we again see that, although significantly lower than in decodable neurons (Wilcoxon rank sum, *p <* 10*^−^*^16^), non-decodable neurons contain a non-zero amount of information about active-empty grasp distinctions. The presence of this information, along with high fidelity decoding from complete population activity in M1, indicates robust encoding of active grasp maintenance at the population level, concentrated in decodable sub-populations but also spread across non-decodable cells in M1.

To further examine the population structure of these responses and understand what aspects of population activity may be supporting strong decoding performance, we analyzed the distribution of PETHs across neurons to compare the relative activation sequence during active and empty grasp trials (Figure S1B). We quantified timing differences at the single-cell level by comparing each neuron’s PETH to itself using random half-splits of trials for both within- and cross-category comparisons. Within-category correlations, comparing empty to empty or active to active trials, were significant higher than cross-category correlations, although the effect size was modest (Figure S1C; paired t-test: effect size = 0.050, p = 2.8 *∗* 10*^−^*^8^). This indicates that despite similar behavioral trial timing and gross kinematic structure (Figure 1F), neurons exhibit small but reliable shifts in activation timing between active and empty grasp trials. To ensure that this result was not an artifact of averaging trials, we repeated the analysis using all pairwise combinations of individual trials and then averaged correlations within each cell. This analysis produced a similar result: within-category correlations were modestly but significantly higher than cross-category correlations (Figure S1D; paired t-test: effect size = 0.0018, p = 7.45 *∗* 10*^−^*^8^). Although these shifts in activation timing may not be large enough to create distinctions at the single cell level, with relatively few neurons exhibiting time-dependent decodable activity (Figures 3C-D), even subtle changes in activation timing can impact the relative timing relationships between neurons to create larger population- or network-level category distinctions. Together with varied single cell representations and strong population decoding accuracy, these findings indicate that individual neurons and populations in M1 represent information about active grasp–beyond that contained in gross or fine kinematics–through firing rate modulation and shifts in activation timing.

### Functional networks are distinct during active and empty grasp

Because reliable shifts in activation timing could alter the correlation structures among simultaneously active neurons, we next examined whether active grasp maintenance was associated with changes in functional network structure. This analysis allowed us to test whether active and empty grasps differed not only in the timing of individual neurons, but also in the broader pattern of pairwise correlations across the recorded population. To quantify these relationships, we constructed condition-specific functional connectivity matrices for each mouse. For each condition, active grasp or empty grasp, we generated an NxN matrix, in which entry (*i, j*) in the matrix *k* was the zero-lag Pearson correlation between simultaneously recorded neurons *i* and *j* over all trials and time points of condition *k*. These matrices, or functional networks (FNs), are signed, weighted, undirected graphs describing pairwise relationships, or functional connections (FCs), among simultaneously recorded neurons during a given condition. The sign of each functional connection indicates whether the correlation was positive or negative, whereas the magnitude indicates the statistical reliability of the correlation. To identify significant functional connections, we compared each observed Pearson correlation to a null distribution generated for the same neuron pair during the same condition using rate- and temporal-dynamics-matched null models for each condition. A functional connection between neurons *(i, j)* during condition *k* was retained if the magnitude of its correlation exceeded the corresponding null distribution with a p-value falling below a 0.05 significance threshold after Bonferroni correction for the number of pairwise interactions per mouse. Connections that did not pass this significance threshold for condition *k* were considered null and set to zero in FN*_k_*. These null models preserve both the average firing rate and carry PETHs for each neuron, so significant functional connections are those whose magnitudes are greater than expected by both rate and carry-related timing dynamics.

Using this metric, we found that decodable neurons were more highly connected in both functional networks, having a higher average number of non-zero functional connections per neuron across either network (121.00*±*0.96) compared to non-decodable cells (115.15*±*0.95). We also found that the active grasp FN was sparser than the empty grasp FN. Active grasp networks contained non-zero connections in 28.17*±*0.45% of all possible interactions, while empty grasp networks contained non-zero connections in 30.27*±*0.72% of all possible interactions (example functional networks in Figure S2A). Although active grasp networks were sparser, the pairwise relationships that remained were stronger than those in empty grasp networks. Active grasp networks had higher magnitude FCs than empty grasp networks for both positively and negatively signed connections (median active grasp FC magnitude = 0.128; median empty grasp FC magnitude = 0.101; Wilcoxon rank-sum, *p <* 10*^−^*^16^; Figures 4A, S2B-C). Comparison with rate- and spatiotemporal activity-matched null models indicated that FC magnitude differences between conditions could not be explained by firing rate differences alone (Figure S2D; *p <* 10*^−^*^16^). This shift in FC magnitude, unexplained by changes in single cell properties like rate and temporal patterning, indicates that in addition to single cells and populations, functional networks distinguish between active and empty grasp.

**Figure 4.**
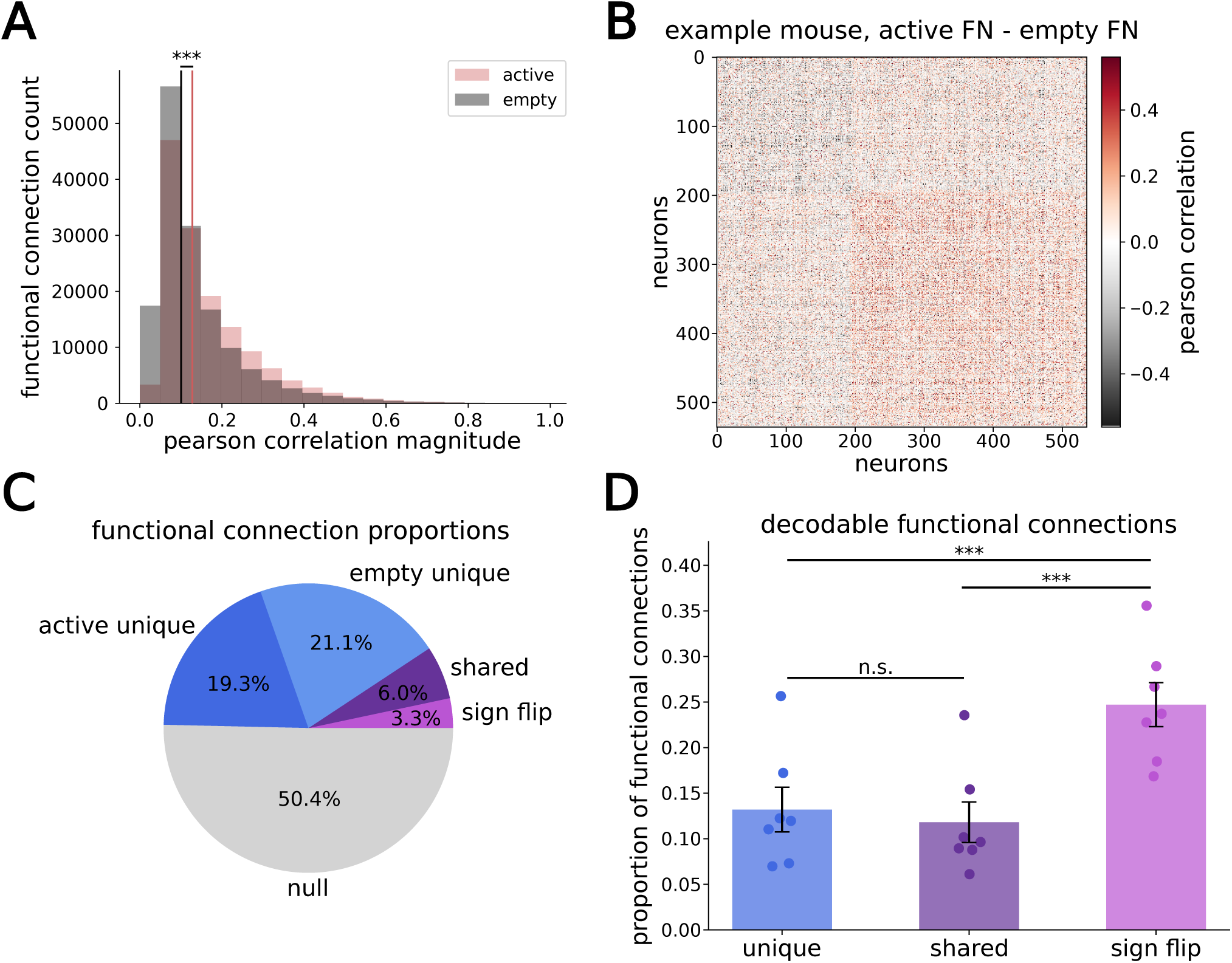
Functional networks are distinct during active and empty grasp. A) Histogram of correlation magnitudes in non-zero active and empty grasp functional networks. Vertical black line indicates median non-null empty grasp functional connection magnitude. Vertical red line indicates median non-null active grasp magnitude. Active grasp functional connections were higher in magnitude than empty grasp functional connections (active median FC magnitude = 0.128, empty median FC magnitude = 0.101, *p <* 10*^−^*^16^ by Wilcoxon rank-sum; n=142190 active functional connections; n=151251 empty functional connections). B) Difference between active and empty grasp functional networks, example mouse. Sorted by multi-resolution consensus clustering ^26,27^. C) Pie chart of functional connections by type. D) Proportion of decodable functional connections by type. Sign flip had a higher proportion of decodable functional connections than shared (*p* = 1.6 *∗* 10*^−^*^5^ by one-sided paired t-test by mice) or unique functional connections (*p* = 2.4 *∗* 10*^−^*^5^ by one-sided paired t-test by mice). Unique and shared functional connections had similar decodable proportions (*p* = 0.070 by two-sided paired t-test by mice; n=7 mice). *** *p <* 0.001, n.s. *p >* 0.05. Also see Figure S2.

Our functional networks were defined by connectivity–which FCs were present or null–and FC signs, in addition to FC magnitudes. Visual examination of functional networks revealed that the connectivity and sign of FCs may also be differentiating between active and empty grasp. When sorting networks by multi-resolution consensus clustering ^26,27^ on the difference between active and empty FNs (Figure 4B, Figure S2A), we saw two clear blocks emerge: one set of neurons with strong functional connections during empty grasp (gray block, upper left, Figure 4B; left, Figure S2A) and a second with strong functional connections during active grasp (red block, bottom right, Figure 4B; right, Figure S2A). This hinted that large sets of functional connections may be condition-specific. Indeed, among FCs present in at least one condition, most were condition-specific, or ‘unique’ (38.9%, n=95943 unique to active grasp; 42.5%, n=105004 unique to empty grasp). A smaller fraction was shared between both conditions (’shared’ FCs, 12.1%, n=29666), and the smallest portion was present in both conditions but with opposite sign (’sign-flip’ FCs, 6.6%, n=16581) (Figure 4C).

Although FN connectivity clearly changes between conditions, since functional networks are calculated using the Pearson correlation, which averages across all trials of a given condition, the presence of these unique and sign flip functional connections does not necessarily mean that the network can distinguish between active and empty grasp on a trial-by-trial basis. To test if FCs can distinguish active from empty grasp in individual trials, we trained SVM decoders to predict trial category, active or empty grasp, from time-varying functional connection sign and magnitude, defined as the edge time series ^28^ between individual pairs of neurons. Across all possible neuron pairs, 14% of functional connections were decodable, defined by cross-validated accuracy and 95% confidence interval above chance. Decodable connections included all FC categories: shared, unique, and sign-flips. Sign-flip FCs were enriched for decodability (24.7% (*±*2.4% of sign-flip FCs were decodable; paired t-test, sign flip vs shared: *p* = 1.6 *∗* 10*^−^*^5^; sign flip vs unique: *p* = 2.4 *∗* 10*^−^*^5^). Shared and unique functional connections showed similar decodability rates, with 11.8% (*±*2.2%) of shared FCs and 13.2% (*±*2.4%) of unique FCs being decodable (paired t-test, *p* = 0.070; Figure 4D). Thus, even among connections that were present with the same sign in both active and empty grasp networks, differences in connection magnitude could distinguish the two conditions in individual trials.

To test whether condition-specific functional connectivity was related to single-cell representations, and to provide an independent predictive validation of the active–empty distinctions identified by decoding, we constructed a family of linear encoding models to predict the activity of individual neurons during active and empty grasp trials. Each model included detailed kinematic terms and a coupling term derived from the activity of the other simultaneously recorded neurons (Figure 5A). This coupling term represented the inferred network input to the predicted cell, computed as the weighted activity of all other cells. Weights were derived from functional networks calculated only on the training data for the model. Because the functional networks were undirected, we treated each significant pairwise correlation as a potential source of input to the predicted neuron for the purposes of the encoding model. We excluded mouse 2 from these analyses because the relatively small number of recorded neurons (n=154 cells) did not support reliable estimation of network structure or coupling effects.

**Figure 5.**
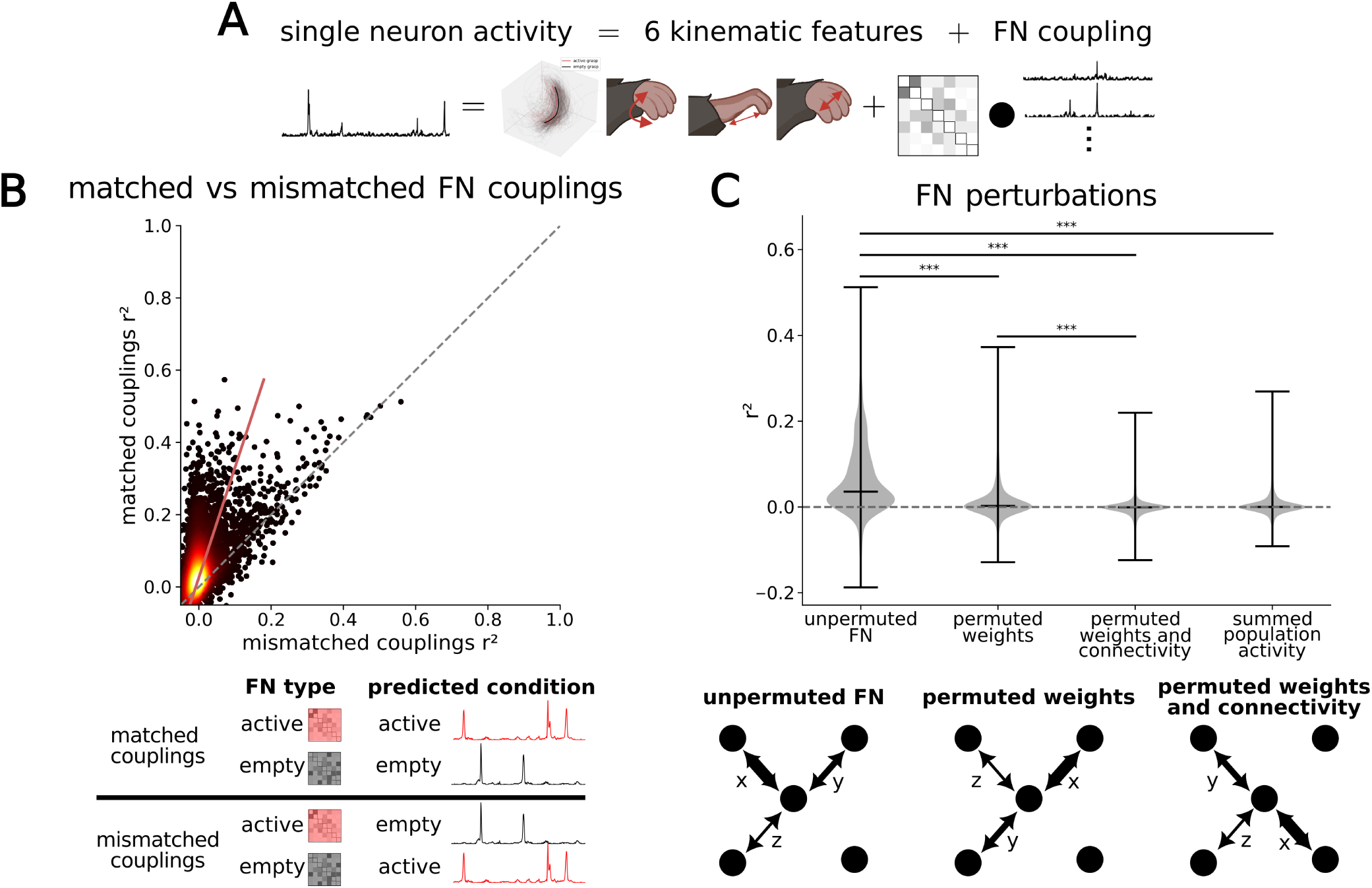
Matched-condition functional connectivity more accurately predicts neural encoding. A) Schematic of encoding model. B) (top) Scatter plot of matched-condition coupling cross-validated *r*^2^ vs mismatched-condition coupling cross-validated *r*^2^ Color indicates density. Red line is line of best fit. (n=4664 neuron-conditions). (bottom) Graphic of matched and mismatched couplings. C) (top) Violin plot of cross-validated *r*^2^ values for functional network perturbation models (n=4664 neuron-conditions). Matched condition model had higher *r*^2^ than permuted weights (*p <* 10*^−^*^16^ by one-sided paired t-test by neuron-conditions), permuted weights and connectivity (*p <* 10*^−^*^16^ by one-sided paired t-test by neuron-conditions), and summed population activity models (*p <* 10*^−^*^16^ by one-sided paired t-test by neuron-conditions; n=4664 neuron-conditions). (bottom) Schematic of functional network perturbations. *** *p <* 0.001. Also see Figure S3.

Because the coupling term was computed from recorded population activity in the training set before fitting the encoding model, rather than learned post-hoc by assigning independent fitted weights to each predictor neuron, it provided a fixed FN-derived estimate of inferred network input. This allowed us to assess the contribution of the functional network as a structured population-level feature, rather than the contribution of individual neurons optimized for model performance. It also allowed us to compare encoding models that differed only in the functional network used to construct the coupling term.

Matched-condition coupling models used the FN computed from training trials of the same condition as the activity being predicted, such as using the active FN to model active grasp activity or the empty FN to model empty grasp activity. Mismatched-condition coupling models used the FN from the opposite condition, such as using the active FN to model empty grasp activity or the empty FN to model active grasp activity (Figure 5B, bottom). Matched condition coupling encoding models performed above chance and explained significantly more variance than kinematics-only models (paired t-test, *p <* 10*^−^*^16^, Figure S3A). In contrast, mismatched condition coupling models performed significantly worse than matched condition coupling models, and closer to the level of the kinematics-only models (paired t-test, *p <* 10*^−^*^16^, Figure 5B). Thus, functional networks from the opposite trial type provided little predictive power for individual neuronal activity. As both sets of models contain the same kinematic features, these results indicate that active and empty grasp functional networks differ in a way that is relevant to single-cell representations and cannot be explained by linear encoding of kinematic differences between the two conditions.

To test whether this condition-specificity could be captured by a linear combination of the condition-unique and shared correlations, we designed a combined-condition encoding model that predicted neural activity across both active and empty grasp trials (see Methods for details). Despite being fit with larger training sets, the combined model did not outperform the matched-coupling condition-specific encoding models (paired t-test, *p* = 2.87 *∗* 10*^−^*^8^ Figure S3B). This indicates that the impact of active-empty FN differences on encoding cannot be fully recapitulated by combining condition-unique and shared coupling terms within a single model. Thus, although shared functional connections contribute to encoding performance, the overall network structure changes sufficiently between trial types that condition-specific coupling provides a better account of single-neuron activity.

To verify that matched, condition-specific encoding performance relied on condition-specific connectivity, rather than merely the aggregate population activity or condition-specific shifts in mean FC magnitude, we performed a series of FN perturbations. We first compared the matched FN-coupling model with an aggregate population model, in which the coupling term was replaced by the summed activity of all simultaneously recorded neurons, excluding the neuron being predicted. This summed-population model performed poorly, with prediction accuracy near chance and significantly lower than the matched FN-coupling model (Figure 5C, paired t-test, *p <* 10*^−^*^16^). We then tested whether the specific structure of the functional network was necessary by permuting the FN weights and connectivity. In one control, FC weights were randomly re-assigned across all possible neuron pairs, disrupting both functional network topology and the magnitude of connections. This model performed at chance and significantly worse than the matched FN-coupling model (Figure 5C, paired t-test, *p <* 10*^−^*^16^). In a second control, weights were permuted only among existing FCs, preserving network topology while disrupting the magnitude and sign assigned to each coupling. This model performed slightly above chance, but still significantly lower than the unpermuted matched FN-coupling model (Figure 5C, paired t-test *p <* 10*^−^*^16^). Each control model also performed worse than the mismatched coupling model (paired t-tests, all *p <* 10*^−^*^16^), indicating that the limited shared structure between active and empty grasp networks supports more predictive power than would be expected if the network were entirely unrelated across conditions. Together, these controls show that encoding performance depends on the specific FN-derived coupling structure, rather than simply on summed population activity or arbitrary coupling among recorded neurons. Because each model included the same condition-matched kinematic features, the selective improvement produced by condition-specific FC weights and connectivity indicates that active and empty grasp functional network structure contributes to single-neuron encoding beyond what can be explained by differences in kinematics alone.

### Active grasp representations become more stable and distinct from empty grasps over learning

After establishing that active and empty grasps are represented in M1 at the single cell, population, and functional network levels during the grasp refinement period, we sought to understand if this representation is stable or emerges over learning. Examining within- and cross-day representational stability and single cell discriminability, we found that changes in the neural representations of active and empty grasp reflected behavioral changes, with active grasp representations stabilizing and differentiating from empty grasp representations over the grasp refinement window.

To examine within-day representational stability across learning, we repeatedly split active and empty grasp trials into random halves and calculated correlations between the resulting peri-event time histograms (PETHs) for each cell. Higher correlations indicate greater within-day consistency across trials. Over learning, within-day correlations of active grasp PETHs increased and plateaued at the active-empty crossover (Figure 6A). This increase indicated more consistent neural representations of carries with active grasp. In contrast, correlations among empty grasp PETHs became persistently lower than active grasp correlations at the crossover point (Figure 6A). Because empty grasps became less frequent over learning, this decrease could, in principle, reflect reduced reliability of PETH estimates from fewer trials rather than a change in the underlying neural representation. However, the difference between active and empty grasp within-day correlations persisted after controlling for trial count (Figure S4A). In addition, after controlling for trial count, cross-day PETH correlations were consistently higher for active grasp than for empty grasp, indicating greater cross-day stability of active grasp representations (Figure S4B).

**Figure 6.**
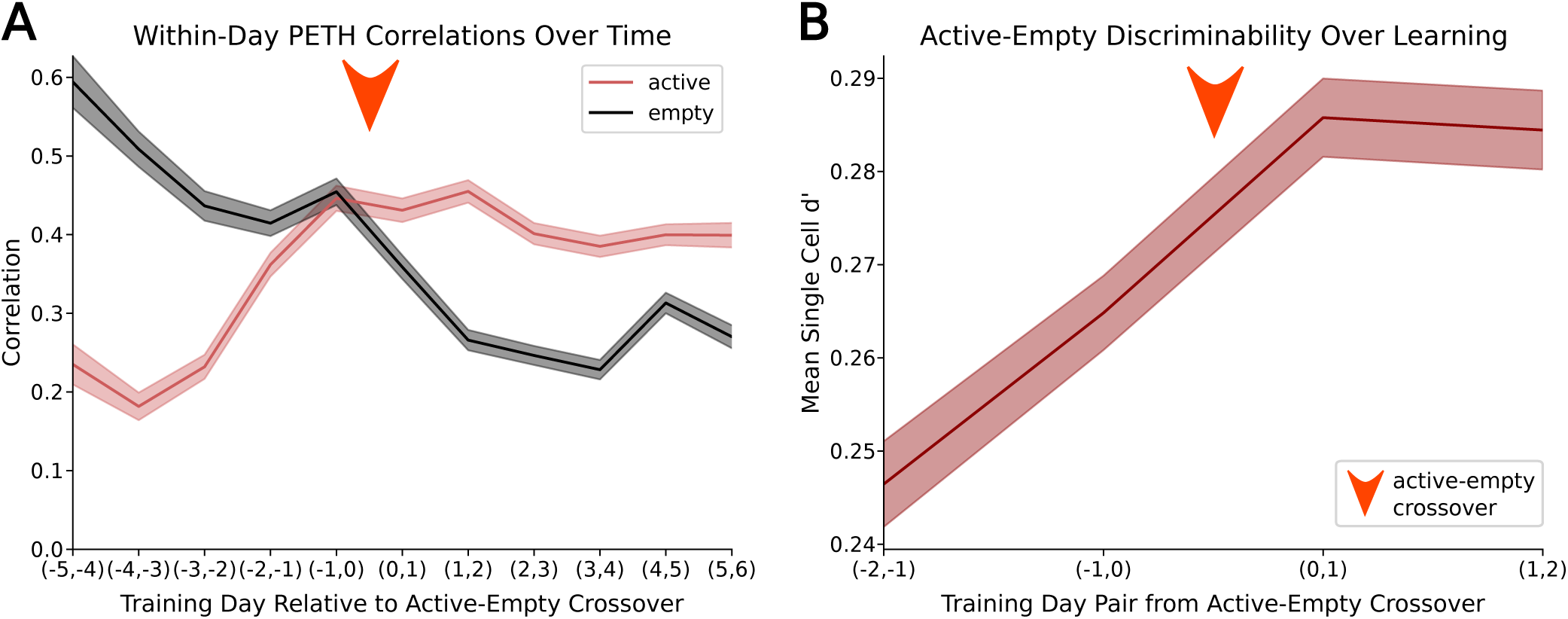
Active grasp representations become more stable and distinct from empty grasps over learning. **A)** Within-day correlation of PETHs by pair of days relative to active-empty crossover. Lines are average across cells, shaded areas are standard error across cells. Orange arrow indicates crossover point. Number of cells aligned across pairs of days varies per pair of days. (n=3118 *±* 317 neurons (mean*±*standard error; range: 889-4241 neurons) B) Discriminability between active and empty grasps by pair of days relative to active-empty crossover, line is population mean d’ across all cells, shaded region is standard error across cells. Orange arrow indicates crossover point. (n=2378 neurons). Also see Figure S4.

Discriminability between active and empty grasp based on single cell activity also increased after the active-empty crossover. Tracking the same cells across all days, mean single cell d’ was significantly higher after the active-empty crossover than before (paired t-test, *p* = 2.84 *∗* 10*^−^*^11^, Figure 6B). These changes across the grasp refinement window indicated M1 increasingly attuning to active grasp as a distinct sensorimotor state from empty grasp and that distinction increasing in stability over learning.

## DISCUSSION

Here, we show that M1 activity distinguishes active grasp maintenance from kinematically similar empty grasp carries, implicating M1 in encoding sensory information in addition to motor information. Active and empty grasp representations were distinct at the level of single-neurons, populations, and functional network structure. These distinctions became more stable and increasingly separable as animals learned to prefer active grasps over empty grasps in a self-paced reach-grasp-carry task. These results suggest that M1 may be involved in sensorimotor integration during behavior that requires it. Additionally, learning-related changes in M1 are not restricted to those underlying refined movement kinematics. Rather, our findings suggest that M1 activity is reorganized during learning to support the tactile-motor coordination required during an actively maintained grasp.

These insights were enabled by a new naturalistic behavioral framework in which mice determined the timing and structure of their own reach-grasp-carry attempts. Because animals could continue into the carry phase after failing to secure the pellet, the task generated empty grasps as naturally occurring ‘catch trials’ in the process of learning. These movements had strong kinematic similarities to active grasp trials. Three dimensional paw and forelimb trajectories were similarly correlated across and within trial categories, and kinematic features supported only modest decoding of active versus empty grasp. Thus, the existence of these trials allowed us to distinguish between neural representations of kinematics, which were largely shared between active and empty grasps, and sensorimotor state, which differed between active grasp–a condition with a pellet and requiring continued incorporation of tactile information to maintain the grasp–and empty grasp–a condition lacking the presence of the pellet entirely.

Distinctions between representations of active and empty grasp also cannot be attributed purely to reward anticipation, successful trial completion, or pellet consumption, which are all known to modulate M1 activity ^22,29–31^. To explicitly exclude any instances of sucrose consumption, we considered a limited set of trial time points that did not include the transition between carry movements and eating. Additionally, the separation between active and empty grasp is not perfectly predictive of reward. While empty grasps could not lead to reward because the pellet had not been secured, active grasps also did not always result in pellet consumption because mice could still drop the pellet as they began attempting to eat it. Furthermore, mice frequently made mouth-directed movements and licked at their paws following empty grasps–behavior consistent with reward expectation despite the lack of pellet presence. These observations argue against a simple account in which active-empty representational differences reflect only task success or expected reward. The most parsimonious interpretation is that the distinction in M1 activity reflects whether the paw is actively engaged with and maintaining control of the pellet during carry, potentially through incorporation of tactile, proprioceptive, grip force, and other signals associated with object contact and grasp maintenance.

A growing body of work supports the view that motor and somatosensory cortices operate as an integrated sensorimotor system rather than as discrete functional units. Sensory and motor cortical areas are highly interconnected ^32–34^, providing an anatomical substrate for rapid communication between regions. Consistent with this organization, sensorimotor information is distributed across both primary somatosensory cortex (S1) and M1^35,36^. In mice, detailed kinematic representations can be found in S1 in addition to M1^37^, paw photostimulation evokes M1 responses ^12^, and proprioceptive signals have been located in M1^38,39^. In primates, both direct tactile stimulation of the hand and the expectation of tactile signals evoke M1 responses and can alter M1 population dynamics ^13–16^. Our findings extend this view by showing that, during naturalistic, dexterous behavior, mouse M1 activity distinguishes between kinematically similar but functionally distinct movements with or without a grasped object. These distinctions, unexplained by subtle kinematic differences between trial categories, implicate M1 in encoding the tactile or proprioceptive signals supporting a successfully maintained grasp. Although we do not directly identify the sensory signals that underlie this differentiation, the existence of a robust distinction between active and empty grasp representations supports a view of M1 as part of a sensorimotor circuit that represents behaviorally relevant variables associated with maintaining an object-related grip.

Our task design also allowed us to examine changes in neural activity linked to grasp-specific learning. Because training sessions lasted a fixed amount of time, mice faced a natural trade-off between time spent completing unrewardable empty grasps and the number of pellet presentations they could attempt. Thus, animals could increase the number of rewards without requiring a corresponding increase in success rate by learning to perform fewer empty grasps. The grasp refinement window–a period where animals learned to preferentially abort trials if the specific sensorimotor state associated with an active grasp was not achieved–is unique to a self-paced task design. When aligning to each animal’s specific refinement window, we found that representations of active grasp became more stable and distinct from representations of empty grasp over time, mirroring behavioral changes. Although these data do not establish whether this neural refinement is required for behavioral improvement, the alignment between the active-empty crossover, increased active grasp representational stability, and greater active-empty discriminability suggests that learning refines M1 activity around the sensorimotor distinctions between empty grasp and active grasp maintenance.

Maintaining an object during carry is not reducible to controlling gross forelimb trajectory alone. It requires fine digit control, appropriate exertion of grip force, and sensorimotor monitoring of the paw-object interaction. The active-empty grasp contrast isolates a behaviorally meaningful M1 state associated with grasp maintenance, characterized by single cell modulation, population-level decodability, and functional network changes, that is not well explained by measured kinematics alone. The stabilization and increased discriminability of this state paralleled the reduction in empty grasps during training, suggesting that learning increasingly structures M1 population activity around the demands of maintaining an object-related grip. These findings show that, during naturalistic skilled dexterous behavior, M1 activity differs according to whether a movement is coupled to an actively maintained grasp. This implicates M1 in sensorimotor, rather than purely motor, computations. More broadly, the results emphasize that successful object manipulation depends not only on generating appropriate movements, but also on integrating motor output with information about whether control of the object is maintained.

## RESOURCE AVAILABILITY

### Lead Contact

Further information and requests for resources should be directed to and will be fulfilled by the lead contact, Jason N. MacLean.

### Materials availability

This study did not generate new, unique reagents. Viral vector and experimental materials used in this study are commercially available. Additional information regarding experimental materials is available from the lead contact upon request.

### Data and code availability

All data reported in this paper will be shared by the lead contact upon request. Custom Python analysis code has been deposited at ithub.com/MacLean-Lab-UChicago/published-code/tree/papers-and-preprints/Wiener-et-al and is be publicly available as of the date of publication. Any additional information required to reanalyze the data reported in this work is available from the lead contact upon request.

## AUTHOR CONTRIBUTIONS

Conceptualization, E.T.W. and J.N.M.; Data Acquisition, E.T.W. and G.W.F.; Analysis, E.T.W.; Funding Acquisition, G.W.F. and J.N.M.; Investigation, E.T.W. and G.W.F.; Methodology, E.T.W. and G.W.F.; Supervision, J.N.M.; Visualization, E.T.W.; Writing - original draft, E.T.W. and J.N.M.; Writing - review and editing, E.T.W., G.W.F., and J.N.M.

## ACKNOWLEDGMENTS

The authors would like to thank Harold Rockwell, Sunnie Hong, and Maria Pope for useful discussions and feedback on the manuscript. This work was supported by National Science Foundation Graduate Research Fellowship Program (NSF GRFP) fellowship DGE-1746045 (G.W.F.) and National Institutes of Health grant R01-NS094184 (J.N.M). Any opinions, findings, and conclusions or recommendations expressed in this material are those of the authors and do not necessarily reflect the views of the National Science Foundation.

## METHODS

### Mice

All experimental procedures were approved by the University of Chicago Institutional Animal Care and Use Committee and were performed in compliance with the Guide for the Care and Use of Laboratory Animals. Data were collected from C57BL/6J mice of either sex (n = 3 female, n = 4 male; Jax strain 007914 bred with either Jax strain 017320 or Jax strain 013044). The lines were originally obtained from Jackson Labs and maintained through in-house breeding. Mice were 11-14 weeks old at the start of training.

### Surgical Procedures

Animals underwent two surgical procedures: AAV-GCaMP6f injections and cranial window and headplate implantation. For all surgeries, anesthesia was induced with isoflurane (induction at 3%, maintenance at 1–1.5%). During AAV injection, three small burr-holes were drilled to allow insertion of a micropipette and injection of virus. Injections were made at two depths (400 and 200 µm below the surface) and in three locations targeting caudal forelimb area (0.25mm anterior and 1.25mm lateral to bregma, 0.25mm anterior and 1.75mm lateral to bregma, and 0mm anterior and 0.5mm lateral to bregma). After injections, the the skin was sutured, and animals recovered for a minimum of seven days before cranial window implantation. During cranial window implantation, a 3.5mm diameter cranial window was implanted above the injection sites and cemented in place with dental cement alongside a custom titanium headbar. Animals again recovered for a minimum of seven days before the start of food restriction and habituation.

### Behavioral Training

#### Food restriction, habituation, and shaping

At least one week after headplate and cranial window placement, mice began food restriction, in which they received a limited amount of pellet food, titrated to maintain their weight between 85% and 90% of their baseline weight. Food restriction and habituation began at the same time. Mice were single housed throughout food restriction, habituation, and training. Mice were habituated to the researcher and to head fixation over at least five days, with head-fixation duration gradually increased before shaping began. During habituation, mice were briefly head-fixed in the two-photon microscope to assess GCaMP6f expression. Only mice with sufficient GCaMP6f expression proceeded to behavioral training.

During shaping, mice were head-fixed inside the 2-photon microscopy rig. On the first day, sucrose pellets were presented on rotating pedestals positioned at licking height. On the second day, pellets were placed in a trough first positioned at licking height, then progressively lowered to a height at which the pellet could be reached with the contralateral forelimb (relative to recording location), but not licked. If the animal began reaching with its ipsilateral forelimb, the handlebar for the ipsilateral side was progressively raised to block ipsilateral and encourage contralateral reaching. To advance from from shaping to training, mice had to successfully reach for and consume at least 15 pellets during a thirty minute recording session. Shaping was performed daily until this criterion was met. The following day would become the first training day. Mice underwent no more than four days of shaping, although some advanced to training after only two days. Imaging took place every day of shaping.

#### Training

During training, sucrose pellets were presented one at a time on rotating delivery pedestals positioned directly in front of the mouse (head-fixed adaptation of pedestal-based reach-to-grasp tasks in de Laittre and MacLean ^40^ and Becker, Calame et al ^41^). A new pellet was delivered after the previous pellet was removed, either through successful retrieval or by being knocked off the pedestal, and the mouse reset to placement of their paws on the handlebar. If the mouse did not initiate a reach for more than 30 seconds, the pedestal was refreshed with a new pellet. Each session consisted of 12 behavioral segments, each with 2.5 minutes of pellet availability followed by a 1–2 minute break during which no new pellets were delivered. This yielded 30 minutes of recorded behavior and neural activity per day. Imaging was performed on every training day.

### Behavioral video recording and kinematic reconstruction

#### Videography and keypoint extraction

High-speed infrared video of reaching behavior was acquired at 200 Hz using two FLIR Blackfly S 16S2C cameras. One camera was positioned approximately 30° from head-on, offset to the mouse’s right and angled 20° from above, to capture both forelimbs and paws. The second camera was positioned approximately 80° from head-on, also offset to the mouse’s right, to provide a detailed view of the right forelimb. Kinematic features were extracted from the video data using DeepLabCut ^24,25^.

#### 3D keypoint triangulation

A 0-lag, 10th order, 15Hz low pass butterworth filter was applied on all keypoints using the scipy function *filtfilt*. Following filtering, time points with high uncertainty (likelihood*<*0.4) were removed. Linear interpolation using the numpy *interp* function was completed over a maximum of 20 frames of high uncertainty data, whose likelihood was then manually set to *>* 0.4. 3D DeepLabCut was used to triangulate the two sets of keypoint data with a likelihood threshold of 0.4^25^.

#### Calculating kinematic variables of interest

The paw centroid was calculated as the mean of the four keypoints located at the digit 2 metacarpophalangeal (MCP) joint, digit 3 MCP joint, and the radial and ulnar protrusions of the wrist, with their mean position representing the approximate center of the palm. Splay was the distance between the proximal interphalangeal (PIP) joints of digits 2 and 3. Aperture was calculated as the distance between the average of keypoints at digits 2 and 3 PIP joints and the average of keypoints at the radius and ulnar protrusions of the wrist. Orientation was the y component of the normal vector to the plane of the palm (defined by keypoints at the digit 2 MCP joint, digit 3 MCP joint, and average of the two wrist keypoints), with the y position of the wrist subtracted such that all negative orientation values represented palm-down orientations while positive values represented palm-up paw orientations.

#### Segmenting behaviors

The DeepLabCut extracted keypoints were used to segment behavior into reach, grasp, and carry components. Behavior was first segmented into candidate behavioral events based on peaks in the wrist keypoint in the y direction of the most lateral camera using the scipy function *find peaks*. A random forest classifier using the scikit-learn class *RandomForestClassifier* was then trained on a subset of human curated data, which included all 2D keypoints from both camera angles, to classify these candidate events into reach, grasp, carry, fidget (when the animal moved its paw without reaching towards the pellet), non-movement (spurious events, related to keypoint jitter or obscured keypoints), and food manipulation. After the classifier achieved greater than 80% accuracy on the training data, it was used to classify the remaining data, with human spot checks to confirm performance.

For carries, the point of rotation in the carry was used as a marker to kinematically align trials. The rotation was based on orientation (see *Calculated kinematic variables of interest*), and the rotation time was designated as the camera frame during which orientation crossed zero when moving from negative (palm down) to positive (palm up). If this rotation did not occur (e.g. mice either kept the paw oriented away from or towards the face for the entirety of the carry motion), the frame at which the paw centroid reached half of its maximum distance from the pedestal was used. For mice where both measures are present, these two measures were generally within the same 2-3 kinematic frames (less than 15ms). As the paw rotation marked the middle of a carry motion, the carry start time was taken as 125ms preceding the carry rotation time, and the carry end was taken as 125ms following the rotation time. This range was chosen based on observed trials to exclude any time points prior to the initial grasp as well as any time points containing pellet consumption behavior.

#### Categorizing carries

Carries were manually classified into empty grasp, active grasp, and drops by a researcher watching the videos from one camera angle. When classification was uncertain, the same event was reviewed from the second camera view.

#### Drop timing

Carries labeled as drops were manually labeled for the frame in which the drop occurred.

#### Success classification and success rate calculation

For each of the mice in each of the days, all instances of grasp and carry were manually classified as a success (defined as resulting in consumption of the pellet by the end of the behavior) or fail (no pellet consumption). Failures included instances where the pellet was no longer present.

Success rate was calculated for each mouse on each training day as the total number of successes divided by the total number of pellet presentations. To determine the number of pellet presentations, pedestal rotation times were tracked during all recording sessions. For each pedestal rotation time, DLC pellet keypoint likelihoods from both camera angles were used to determine if the pellet was present. As the pedestals were narrow, pellets sometimes fell off the pedestal before the pedestal rotated into place in front of the mouse. A pellet was determined to be present if the pellet likelihood was *>* 0.9 in keypoints from both camera angles for at least 125ms in the time from 50ms preceding to 200ms following the pedestal stopping in front of the mouse. This amount of time was chosen after observing videos to account for spontaneous pellet dislodgement but not animal-related task behavior.

#### Interpolation

For the linear encoding model analysis, kinematic data (200Hz) was interpolated to the time points of the neural data (30Hz) using the numpy *interp* function.

### Two-photon calcium imaging

#### Image acquisition

Two-photon calcium imaging was performed using a Bruker microscope in a single imaging plane at 30 Hz. The field of view (FOV) was approximately 828×828 µm, sampled at 512×512 pixels. Consistent with the behavioral design, imaging was continuous during each 2.5 minute behavioral segment and was paused during breaks. For analysis, all data from the session was concatenated and analyzed together.

#### Locating the same field of view over days

The same FOV was imaged across all shaping and training days for each mouse. Prior to the first day of training, the imaging coordinates were selected to maximize the number of GCaMP-expressing neurons. The average number of neurons in one FOV on a given day was 810 cells (range: 420-1262). A surface blood vessel image was used to identify the XY coordinates, and statically labeled cells served as fiduciary markers for alignment in the same plane on subsequent days. Imaging was performed at an estimated depth of 200–300 µm below the cortical surface.

#### Registering cells across days

Cells were registered across days using CellReg ^42^. For all cross-day analyses, included neurons were restricted to cells detected on all aligned days, yielding a mean of 355 neurons per mouse for most analyses (range: 154–536).

#### Pre-processing of calcium imaging data

Imaging data were preprocessed in Suite2p (v0.14.5) to identify cells, extract spatial footprints, and obtain background-subtracted single-cell fluorescence traces ^43^. Fluorescence traces were then processed with the spike-inference algorithm CASCADE, and the resulting inferred spike counts were used as the neural activity signal for all analyses ^44^.

### Quantification and statistical analysis

#### Peri-event time histograms (PETHs)

PETHs were calculated from neural activity aligned to paw rotation during the carry. For visualization, PETHs are shown from 150 ms before to 400 ms after carry onset. Quantitative analyses were restricted to 0–250 ms after carry onset to avoid activity preceding grasp establishment or following the onset of reward consumption. For population visualizations, each cell’s PETH was normalized within the -150-400ms window.

To compare PETHs within pairs of days and between active and empty grasps, trials were randomly split in half within each carry category. From each trial split, a PETH was calculated for each cell (for a total of 4 PETHs per cell, two per carry category). For within-category comparisons, a Pearson correlation was computed between two PETHs of that condition. For across-category comparisons, a Pearson correlation was computed between one PETH from either condition. Splits were repeated 1000 times, then correlations were averaged per cell and comparison type across repeats.

To compare PETHs across days, each cell’s PETH was calculated separately from all trials of the same type for each day. Pearson correlations were computed between the same cell’s PETHs on all pairs of consecutive days (e.g. training days 1 and 2, then 2 and 3). The mean correlation across cells for each pair of days was used as the measure of cross-day representational similarity.

#### Determination of decodable cells

A linear kernel support vector machine (SVM) was used to predict trial category from single-neuron activity. Two models were trained for each neuron: a single feature model using mean activity across the full trial duration, and an eight feature model, using instantaneous inferred spike count of a given neuron at 8 time points in a trial, which corresponds to approximately 240 ms at our 30 Hz imaging rate. Models were trained using a 90/10 train/test split and downsampling for active/empty class balancing. Neurons were classified as decodable if the 10-fold cross-validated accuracy, including the 95% confidence interval, exceeded chance for either model. Decodable neurons were further classified as time-dependent if only the eight-feature model exceeded chance. They were classified as increased or decreased if they were decodable and showed higher or lower mean activity, respectively, during active grasp trials compared with empty grasp trials.

#### Kinematics decoder

Three linear kernel SVMs were trained to predict trial category from the 3D kinematics in each trial. For the gross kinematic decoder, the predictors were the instantaneous x, y, and z position of the paw (3 kinematic parameters at 8 time points each for a total of 24 features). For the fine kinematic decoder, the predictors were the splay, aperture, and orientation at each timepoint in the trial (3 kinematic parameters at 8 time points each for a total of 24 features). For the full model, all six fine and gross kinematic parameters were used (total of 48 features). Any trials where any of the six kinematic parameters contained missing data (from the paw being obscured relative to the cameras) were excluded. Models were trained using a 90/10 train/test split and downsampling for class balancing. Scikit-learn’s *GridSearchCV* function was used on training data to select between L1 and L2 regularization and fit the regularization parameter for each fold. Accuracies shown are 10-Fold cross-validated accuracies on balanced data.

#### Neural population decoder

A linear kernel SVM with L1 regularization was trained to predict trial category from population activity of each mouse. The predictors were the average firing rate across all time points in the given trial for each simultaneously recorded neuron. Models were trained using a 90/10 train/test split and downsampling for class balancing. Scikit-learn’s *GridSearchCV* function was used on training data to fit the regularization parameter for each fold. Accuracies shown are 10-Fold cross-validated accuracies on balanced data.

For decoder training over days, pairs of days are used to provide a sufficiently high trial count for each model and accuracies shown are the 10-Fold cross-validated accuracies.

#### Mutual Information between Neurons and Carry Type

Using the Java Information Dynamics Toolkit ^45^, we estimated mutual information between single cell neuronal activity and trial type with a continuous-discrete Kraskov estimator ^46^. Estimations were repeated 100 times, then the average of the repeated estimates was taken as the mutual information value.

#### Functional networks calculation and determination of significant correlations

Functional connections were defined as the Pearson correlation between each pair of neurons. To identify correlations that could not be explained trivially by co-varying firing rates, we constructed a firing-rate and trial-timing null model of functional connectivity using weighted random resampling of neural activity. This resampling strictly preserved each neuron’s overall firing rate across trials, and weights were based off of carry PETHs such that resampled data closely approximated trial-averaged time-varying activity dynamics.

We used repeated weighted random sampling with a reservoir, adapted from Algorithm *A-Res* in Efraimidis, P. S., and Spirakis, P. G. (2006) ^47^. In each resample, firing rates were assigned from the neuron’s recorded neural activity in ascending order according to the value of a key *k_i_*for each time point *i* such that time points with the lowest key values were assigned the lowest firing rates. Let *X ∼ U* (0, 1) be a uniform random variable and *P*(*f_j,t_*) be the probability of neuron *j* firing at time *t* during any carry trial, calculated from the carry PETH of that cell. At each time point *i*, the key value is

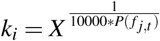

where *t* is the time since the start of the trial for time point *i*. This means that all trials were assigned identical sequences of resampling weights–preserving trial-timing structure–while incorporation of the uniform random variable resulted in a new, semi-random shuffle of firing rates for each null replicate.

Pearson correlations were then calculated between the resampled activity of all neuron pairs. This process was repeated 1000 times to construct a null distribution for each functional connection. Observed pairwise correlations were then compared with the null distribution to identify functional connections that exceeded those expected from shared firing-rate structure alone.

Correlations were considered significant if their magnitude exceeded the null distribution at p*<*0.05 after Bonferroni correction for multiple comparisons (with the number of functional connections per mouse). Non-significant correlations were set to 0.

For visualizations, functional networks were sorted per mouse using multiresolution consensus clustering ^26^ on the difference between the active and empty functional networks, implemented using the Brain Connectivity Toolbox ^27^.

#### Determination of unique and shared correlations

Unique correlations were non-zero (significantly stronger than the null model) in one condition and zero (not significantly stronger than the null model) in the other. Shared correlations were non-zero in both conditions and had matched signs. Sign-flipped correlations were non-zero in both conditions but changed sign, switching between positive and negative correlations across conditions.

#### Determination of decodable functional connections

Similarly to single-neuron decoding, SVMs were trained to distinguish active and empty grasp trials using pairwise functional connectivity. For each neuron pair, models used either the Pearson correlation over the entire trial as a single feature or the functional connection time series (as in Zamani-Esfahlani et al. (2020) ^28^) for each of the eight time points in a trial as an eight-feature representation. Models were trained using a 90/10 train/test split and downsampling for class balancing. A functional connection was considered decodable if the 10-Fold cross-validated accuracy including the 95% confidence interval exceeded chance performance for either model.

#### Condition-Specific Encoding Models

Ridge regression was used to model inferred spike rates. All linear models were fit using scikit-learn’s *Ridge* class. For the full model, the ridge parameter was selected with sckit-learn’s *GridSearchCV* function over a range of ridge parameters between 0 and the total number of time points. All models used 2-Fold cross validation with 10 repeats for a total of 20 cross validation folds with matched train/test splits across models. Time points where any kinematic parameters contained missing data (from the paw being obscured relative to the cameras) were excluded from training and testing datasets. Cross-validated *r*^2^ values are reported as the median across folds. Splits were performed by trial to prevent any train-test data leakage from highly correlated time points within the same trial. Cross validation folds were stratified by the predicted neuron’s firing rate and day labels, ensuring that training and test sets had similar average firing rates and balanced trial distributions across pooled days.

The base model included six kinematic predictors (paw centroid x, y, and z position (*c*(*t*)); splay (*s*(*t*)); aperture (*a*(*t*)); and orientation (*o*(*t*))), along with a coupling term, and a constant to account for average firing rate offset. The coupling term was defined as the linear combination of inferred activity of all neurons, weighted by functional connections to the modeled neuron as calculated from the training data (as in Kotekal and MacLean (2020) ^48^). Because the functional network is undirected, all significant functional connections involving neuron *i* were treated as inputs to neuron *i*. For compact notation, let *r^k^*(*t*) be the vector of inferred firing rates at a given time *t* during the condition *k* and *N* be the total number of neurons in the population. We define 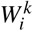 *∈* [−1, 1]*^N^* as the vector of functional weights between neuron *i* and all other neurons in the population:

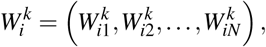

where each element 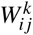 is the Pearson correlation between neuron *i* and neuron *j* during condition *k*, calculated from the training data.

With the coupling and kinematics terms defined, for each neuron i, we constructed a linear model with ridge regularization to model its inferred spiking rate,

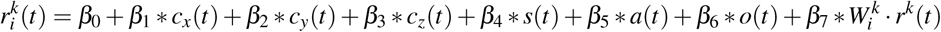

Here, all *β* terms are fitted free parameters. The constant *β*_0_ accounts for differences in average firing rates between neurons. Since the vector of functional weights 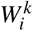 is computed directly from the training data, the vector 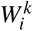 *· r^k^*(*t*) is a feature and not a free parameter. Thus, the full model has eight free parameters.

We fit two condition-specific full models: matched and mismatched couplings. For the matched coupling encoding models, the functional network used to define 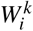 was calculated from the training data within each cross-validation fold. In mismatched-coupling models, 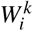 was derived from the functional network calculated using all trials of the opposite category. For example, when predicting active-grasp activity in the mismatched-coupling model, the coupling weights were derived from the empty-grasp functional network. In both models, the kinematic predictors and population activity vector *r*(*t*), which is combined with the functional network weights to define the coupling term, were taken from the same time points as the train and test samples used to predict the target neuron.

All features were normalized by their maximum absolute value in the training data so that each training feature spanned a maximum range of -1 to 1. Test features were normalized using the corresponding set of training set values and therefore were not restricted to -1 to 1. The target neuron’s activity was not normalized in the training or testing datasets.

#### Combined-Condition Encoding Models

To determine how condition-specific functional networks contribute to encoding, we compared the condition-specific full model with a condition-combined model. In the condition-combined model, the coupling term was decomposed into three terms: one for functional connections shared across active and empty grasp trials, and one each for functional connections unique to a condition. Sign-flipped functional connections were not included. The shared-correlation coupling term used weights calculated from both active and empty grasp trials in the training data. In contrast, the condition-unique coupling terms used weights calculated only from the corresponding trial category within the training set. Because this model included more features than the condition-split model, its ridge parameter was selected using *GridSearchCV*, with internal train-test splits within the training data to find an optimal value for the ridge parameter.

For comparisons to the condition-specific encoding models, the condition-specific encoding models’ cross-validated *r*^2^ is averaged between the two conditions. This averaged cross-validated *r*^2^ is then compared to the median *r*^2^ across folds in the combined-condition encoding model for each neuron.

#### Unweighted Input Encoding Models

To determine the extent of the contribution of functional network weights on encoding (rather than just information from other neurons), we compared the matched-condition full model to an unweighted model, which uses the unweighted sum of the activity of all other neurons as the coupling term.

#### Permuted Functional Network Encoding Models

To assess whether encoding depends on the identities of functional connections, rather than only on their weights, we compared the matched-condition full model with permuted functional network models. We used two permutation controls: weight-and-connectivity permutations and weight-only permutations. In the weight-and-connectivity permutation, Pearson correlations calculated from the training data were randomly reassigned to non-diagonal entries of the connectivity matrix, regardless of whether that neuron pair had a significant correlation in the observed network. This permutation disrupted both connection identity and network topology. In the weight-only permutation, the functional network topology was preserved, but the correlation weights, including magnitude and sign, were permuted across existing functional connections. For each model, permutations were repeated 100 times per cross-validation fold. Model performance was averaged across permutations within each fold and then compared with the matched-condition full model.

#### Leave-One-Out Encoding Models

With the exception of the coupling leave-one-out model, equivalent to the kinematics-only model, and the coupling-only model, all leave-one-out variants used a ridge parameter equal to half the number of trials in the training and testing datasets, defined as the number of time points divided by sixteen. This fixed regularization parameter was used to reduce computation time during repeated functional connection count-matched leave-one-out analyses. These analyses were restricted to neurons whose full model, fit with the same ridge parameter, achieved cross-validated *r*^2^ > .05. This ensured that baseline model performance is sufficiently above chance to detect decrements in model performance after removing functional connections from the coupling term.

All leave-one-out models preserved the kinematic predictors and modified only the coupling terms, keeping the total number of features constant.

There were two types of of leave-one-out (LOO) models: complete and count-matched. In complete LOO models, all functional connections of a given type were set to zero when computing the coupling term. Correlation types included shared, matched-condition unique, mismatched-condition unique, sign-flipped, strong magnitude, medium magnitude, and weak magnitude. For magnitude-based LOO models, strong magnitude functional connections were defined as those in the 90th percentile and above by magnitude, medium magnitude functional connections were defined as those between the 25th and 90th percentile by magnitude, and weak magnitude functional connections were defined as those in the 25th percentile and below by magnitude. In count-matched LOO models, a random subset of functional connections from a given type was set to zero, with the subset size matched to the number of functional connections in a smaller functional connection category. Count-matched LOO models were repeated 500 times per fold, and the mean *r*^2^ value across bootstraps is taken as the *r*^2^ value for that fold.

#### Application of Encoding Models on Drop Trials

Encoding models were trained as matched-condition coupling models on a random 70% of empty or active grasp time points for 10 cross-validation repeats (see *Condition-Specific Encoding Models*). These models were then tested on either pre-drop time points (the 125ms preceding the drop) or post-drop time points (the 125ms succeeding the drop), resulting in four model train-test versions per cell (training on active or empty grasp and testing or pre- or post-drop time points). This approach avoided directly estimating functional networks from the limited number of drop time points, which would have been noisy and unreliable. Reported *r*^2^ values are the median across cross-validation repeats.

#### Determination of Drop-Modulated Cells

Drop modulated cells were determined by performing a one way ANOVA between the pre- and post-drop time points (125ms before compared to 125ms after a drop) of the same trials over all aligned days. Because when drops occurred was not consistent across trials, averaging drop-aligned activity across trials should reduce the contribution of stereotyped carry kinematics tied to fixed moments in the movement. This analysis therefore emphasized neural changes associated with pellet loss rather than differences easily attributable to the kinematics of the carry motion itself. A cell was considered modulated if it has p*<*0.05 after Benjamini-Hochberg false discovery rate correction.

## SUPPLEMENTAL INFORMATION

Document S1. Figures S1-S5

Video S1. Active grasp instance, related to Figure 1

Video S2. Empty grasp instance, related to Figure 1

