## Supplemental Information for "Representations of active grasp maintenance emerge during reach-grasp-carry learning in mouse motor cortex"

**Video S1. Active grasp instance.**. Example active grasp trial, extended to show grasp and pellet consumption, slowed to 15% of original speed. Related to Figure 1

**Video S2. Empty grasp instance.**. Example empty grasp trial, extended to show grasp and paw licking behavior, slowed to 15% of original speed. Related to Figure 1

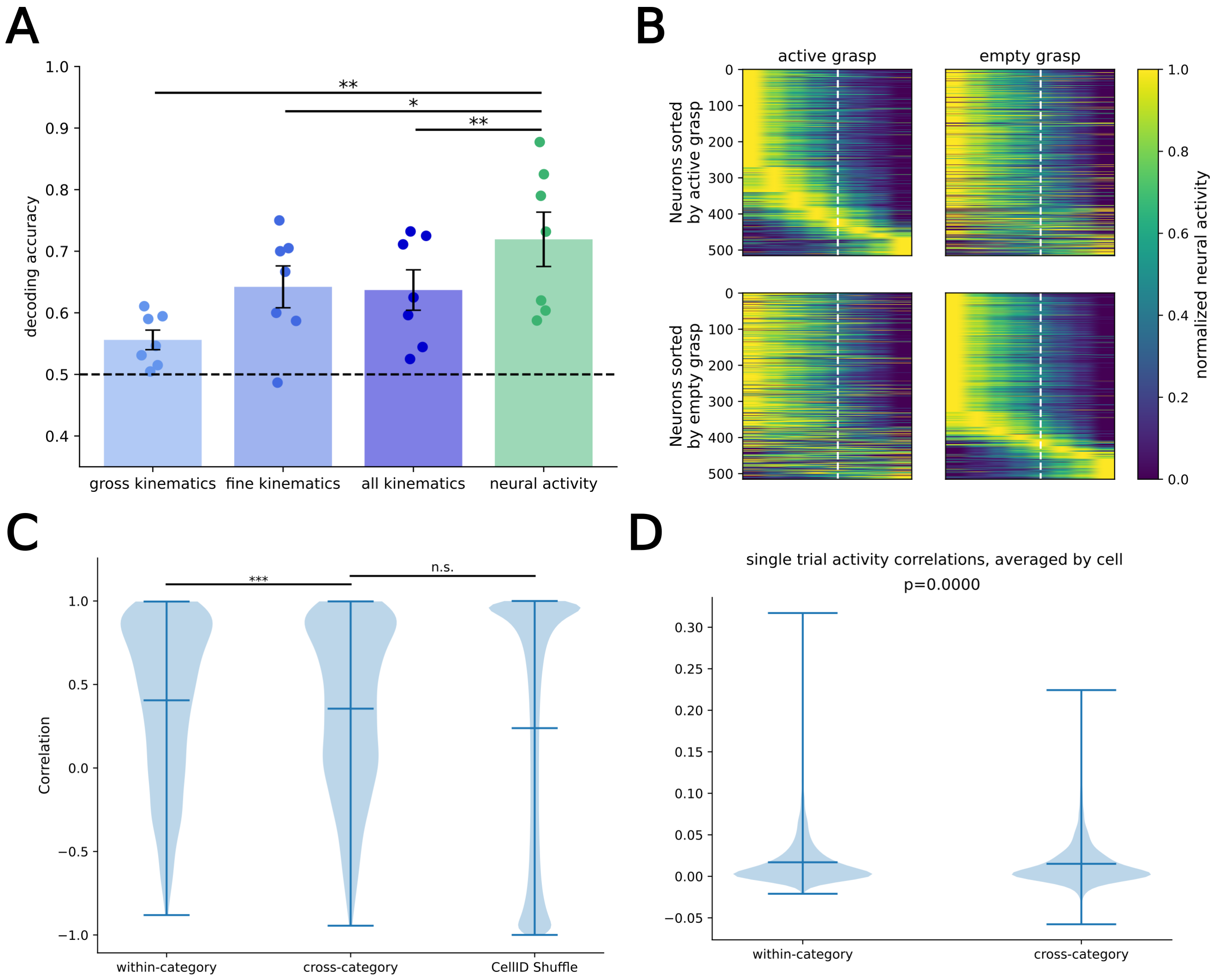


**Figure S1**. Population-level neural activity structures differ between active and empty grasp, related to Figure 3. A) Decoding accuracies for kinematic and neural population models. Accuracy is average across mice, error bars are standard error across mice. Points are individual mouse cross-validated accuracies. Decoding from neural activity outperforms all kinematics ($p = 0.002$ by one-sided paired t-test), fine kinematics ($p = 0.018$ by one-sided paired t-test), and gross kinematics ($p = 0.0012$ by one-sided paired t-test; n=7 mice). B) Example mouse, population PETHs. C) PETH correlations within- and across-categories compared to a CellID shuffled control. Within-category correlations are higher than cross-category correlations (effect size=0.050, $p = 2.8 * 10^{-8}$ by one-sided paired t-test, n=2486 cells). No significant difference between cross-category correlations and CellID shuffle (p=0.14 by Wilcoxon rank-sum, n=1000 CellID shuffle repeats). D) Single trial PETH correlations. Wiithin-category correlations are higher than cross-category correlations (effect size=0.0018, $p = 7.45 * 10^{-08}$ by one-sided paired t-test, n=2486 cells). *** $p < 0.001$, ** $p < 0.01$, * $p < 0.05$, n.s. $p > 0.05$

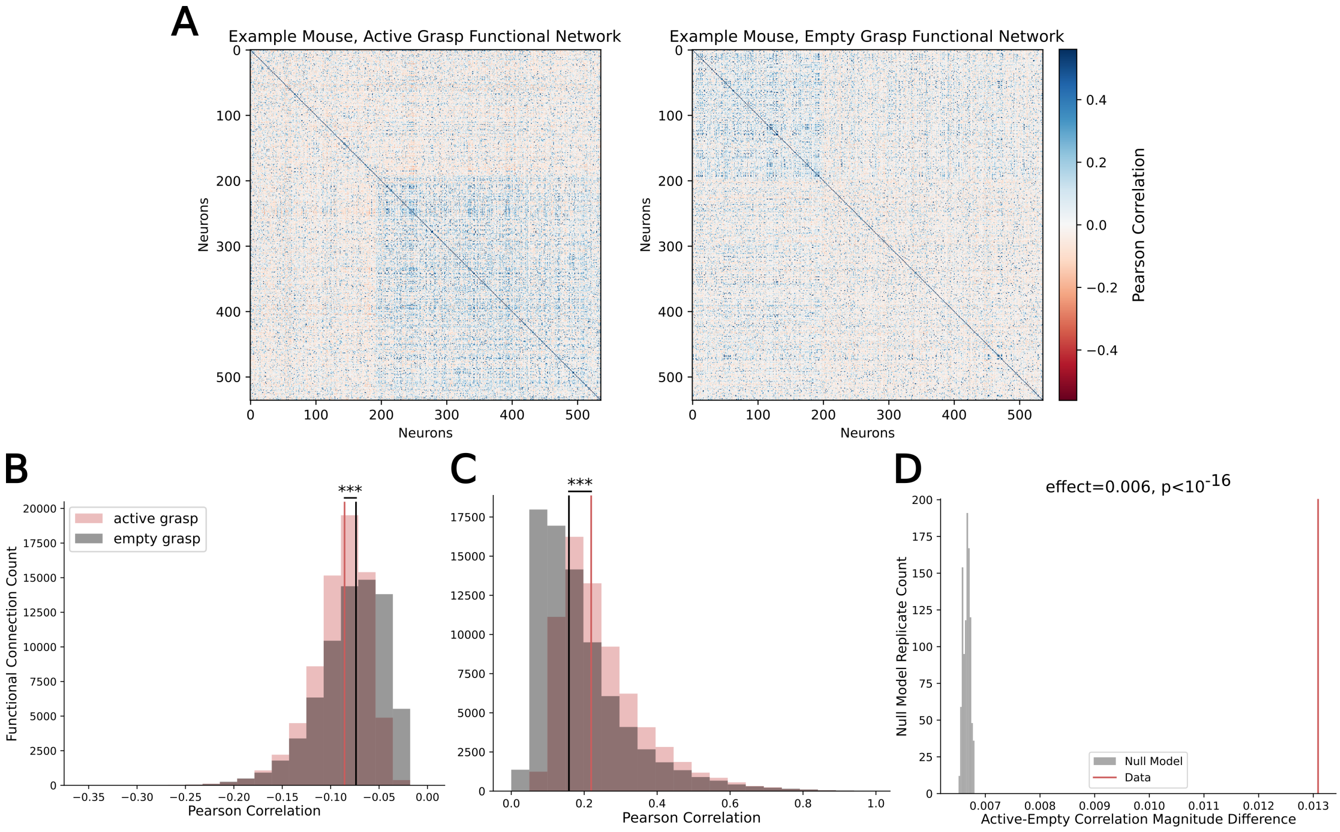


**Figure S2**. Functional networks of active and empty grasp are more distinct than null models, related to Figure 4. A) Example mouse, active (left) and empty (right) functional networks. Sorted using mutliresolution consensus clustering on the difference between active and empty functional networks. B-C) Histogram of active and empty functional connection correlation values. Vertical lines indicate distribution medians. (B) Negative functional connections. Active grasp has higher magnitude (lower signed) correlations than empty grasp ($p < 10^{-16}$ by Wilcoxon rank-sum, n=72636 active grasp FCs; n=72434 empty grasp FCs). (C) Positive functional connections. Active grasp has higher magnitude correlations than empty grasp ($p < 10^{-16}$ by Wilcoxon rank-sum, n=69554 active grasp FCs; n=78817 empty grasp FCs). D) Rate- and spatiotemporal activity-matched null model average magnitude difference between active and empty correlations on different null replicates compared to data. *** $p < 0.001$

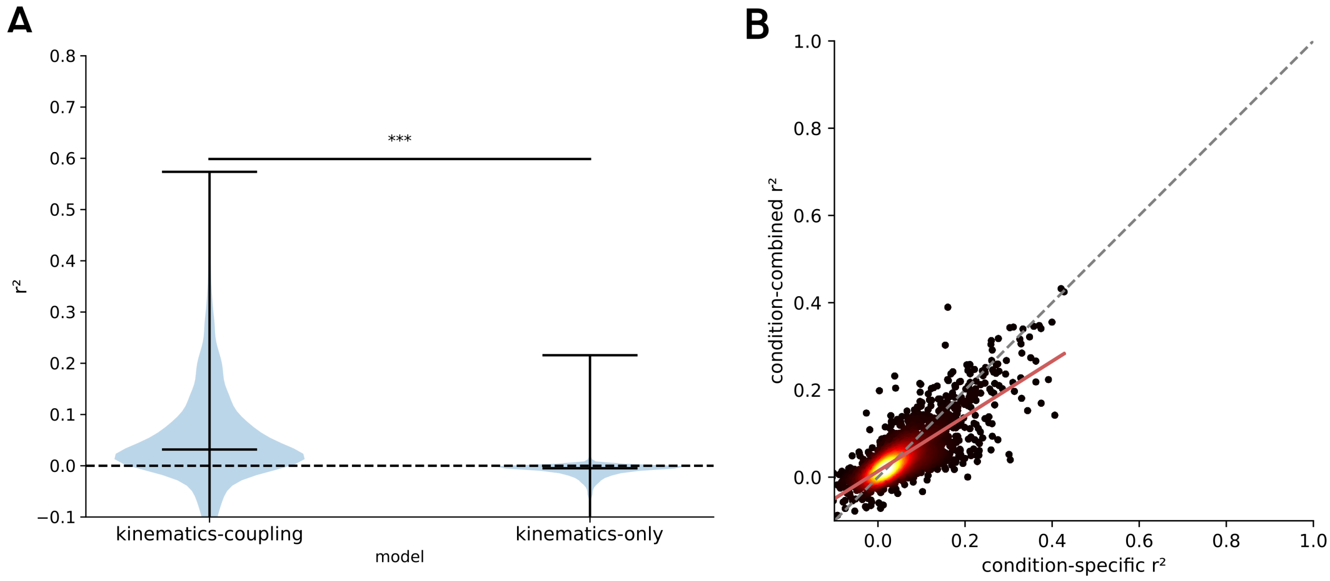


**Figure S3**. Condition-specific encoding models are the most predictive of single cell activity, related to figure 5. A) Kinematic-only encoding model compared to matched-category kinematic and coupling encoding model r2. $p < 10^{-16}$ by one-sided paired t-test (n=4664 neuron-conditions). B) Combined- compared to condition-specific encoding model performance. Colormap indicates density. Pink line is line of best fit. Condition-specific $r^2$ values are higher than condition-combined $r^2$ values ($p = 8.1 * 10^{-8}$ by one-sided paired t-test, n=2378 neurons) *** $p < 0.001$

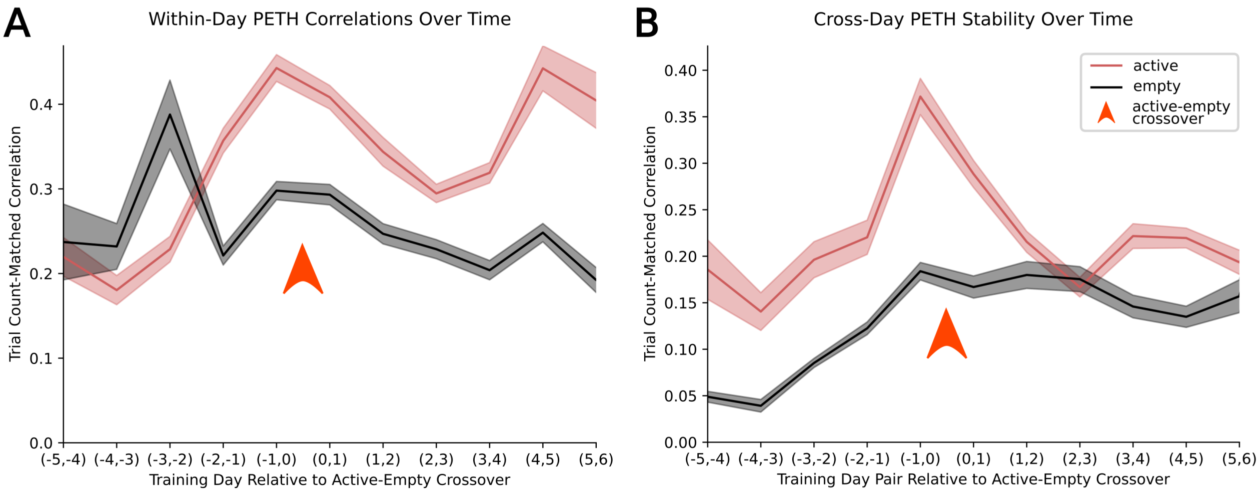


**Figure S4**. Trial count-matched controls show increased active grasp stability, related to figure 6. A) Within-day pair PETH correlations subsampled such that active and empty grasp included the same number of trials per day pair, repeated bootstrap. Lines are average across all cells, shaded areas are standard error across all cells. Orange arrow indicates crossover day. Number of cells aligned across pairs of days varies per pair of days. (n=$3118 \pm 317$ neurons (mean±standard error; range: 889-4241 neurons) (B) Cross-day PETH correlations, subsampled such that each day included the same number of active and empty grasp trials in the PETH, repeated bootstrap. Lines are average across all cells, shaded areas are standard error across all cells. Orange arrow indicates crossover day. Number of cells aligned across pairs of days varies per pair of days. (n=$3118 \pm 317$ neurons (mean±standard error; range: 889-4241 neurons)
